# Longitudinal Decline of Exercise Capacity in Male and Female Mice

**DOI:** 10.1101/2024.07.29.605646

**Authors:** Megan L. Pajski, Rosario Maroto, Chris Byrd, Ted G. Graber

**Author notes:** **Corresponding Author:** Ted G. Graber, PhD., ECU College of Allied Health Sciences, Department of Physical Therapy, Health Sciences Building, Rm 2410 | Mail Stop 668, Greenville, NC 27834.

## Abstract

The population of older adults is exponentially expanding. Alongside aging comes the onset of chronic disease, decline of functional capacity, and reduced quality of life. Thus, this population increase will stress the capacity and financial viability of health and long-term care systems. Developing pre-clinical models for age-related functional decline is imperative to advancing therapies that extend healthspan and prolong independence. Previously in a cross-sectional study, we established a powerful composite scoring system we termed CFAB (comprehensive functional assessment battery). CFAB measures physical function and exercise capacity using well-validated determinants to measure overall motor function, fore-limb strength, four-limb strength/endurance, aerobic capacity, and volitional exercise/activity rate. In the current work, we used CFAB to track cohorts of male and female C57BL/6 mice over the lifespan (measuring CFAB at 6, 12, 18, 24, and 28 months of age). Overall, we found statistically significantly declining function as the mice aged, with some differences between males and females in trajectory and slope. We also determined that body mass changes presented differently between sexes, and tracked body composition (fat percentage, using magnetic resonance imagery) in females. In a subset of mice, we tracked *in vivo* contractile physiology noting declines in plantar flexor maximum isometric torque. In summary, our data suggest that males and females declined at different rates. We confirmed the efficacy of CFAB to track longitudinal changes in exercise capacity and physical fitness in both males and females, further validating the system to track age-related functional decline.

## Introduction

By 2050 the older adult population over the age of 60 is expected to explode to over 2 billion people.(1) Alongside this population growth of older adults will be increasing cases of age-related chronic diseases and syndromes such as dementia, frailty, sarcopenia, cardiovascular disease, kidney disease, diabetes, and many others—including incidence of multimorbidity.(2–7) One common theme of many of these age-associated conditions is declining physical function and eventual loss of independence.(7,8) For example, while estimates of prevalence vary wildly, frailty and sarcopenia respectively affect roughly 11% and 13% of older adults.(3,4) Both frailty and severe sarcopenia, which are often comorbid, are hallmarked by a degeneration of functional capacity leading to decline in ability to perform activities of daily living with eventual loss of independence.(3,4,9) Strategies to preserve physical function may likely rely on multiplex interventions to combat the multiplex etiology of frailty and sarcopenia, with their development requiring validation of reliable preclinical systems to test efficacy.

In previous work we established CFAB (comprehensive functional assessment battery) to measure physical function and exercise capacity in the mouse model using C57BL/6 male mice in a cross-sectional studies with cohorts at adult (6m, months), early older adults (24months, 24m), and older adult ages (28m).(10) CFAB is a composite scoring system consisting of 5 previously well-validated determinants. (10) CFAB has proven to be a valuable resource which we have used to explore the relationship between physical function, aging, and the transcriptome, and to evaluate the efficacy of different exercise paradigms to preserve functional status in both early middle age and older adult male mice.(11,12) In the current work our primary purposes were twofold: 1) to validate CFAB in female mice, and 2) to further demonstrate longitudinal utility over the adult to older adult mouse lifespan. We hypothesized significant (p<0.05) age-related declining function with some sexual dimorphisms in trajectory and slope of decline.

To accomplish our aims, we measured CFAB in female and male mice longitudinally, testing at 6, 12, 18, 24 (25 males), and 28m (29 males). In addition, we tracked body mass, and both fat percentage and plantar flexor torque (in a subset). Importantly we uncovered sexual dimorphisms in the slope of decline and in the way individual CFAB determinants reacted to advancing age— as well as the relationship between body mass and function. Surprisingly, the linear relationship between age and CFAB was stronger in females than males with the slope of decline about 3 times faster. Overall, we successfully demonstrated the ability of CFAB to measure function in both sexes and over the lifespan.

## Methodology

### Mice

All experiments were approved by the ECU Institutional Animal Care and Use Committee and were conducted with animal welfare as the primary consideration. Male (n=20) and female (n=32 at 6m for CFAB baseline, then n=24 for aging) C57BL/6 mice were aged from 5/6m to 28+m, fed/watered ad libitum and housed at ∼73°C on a 12 hr./12 hr. light/dark cycle. The males (C57BL/6JN) were sourced from the NIA Aging Rodent Colony and the females (C57BL/6NCrl) were purchased from Charles River aging colony. See baseline characteristics in **Supplementary Tables S1** and **S2**.

***Study Design:*** See **Figure 1**.

**Figure 1.**
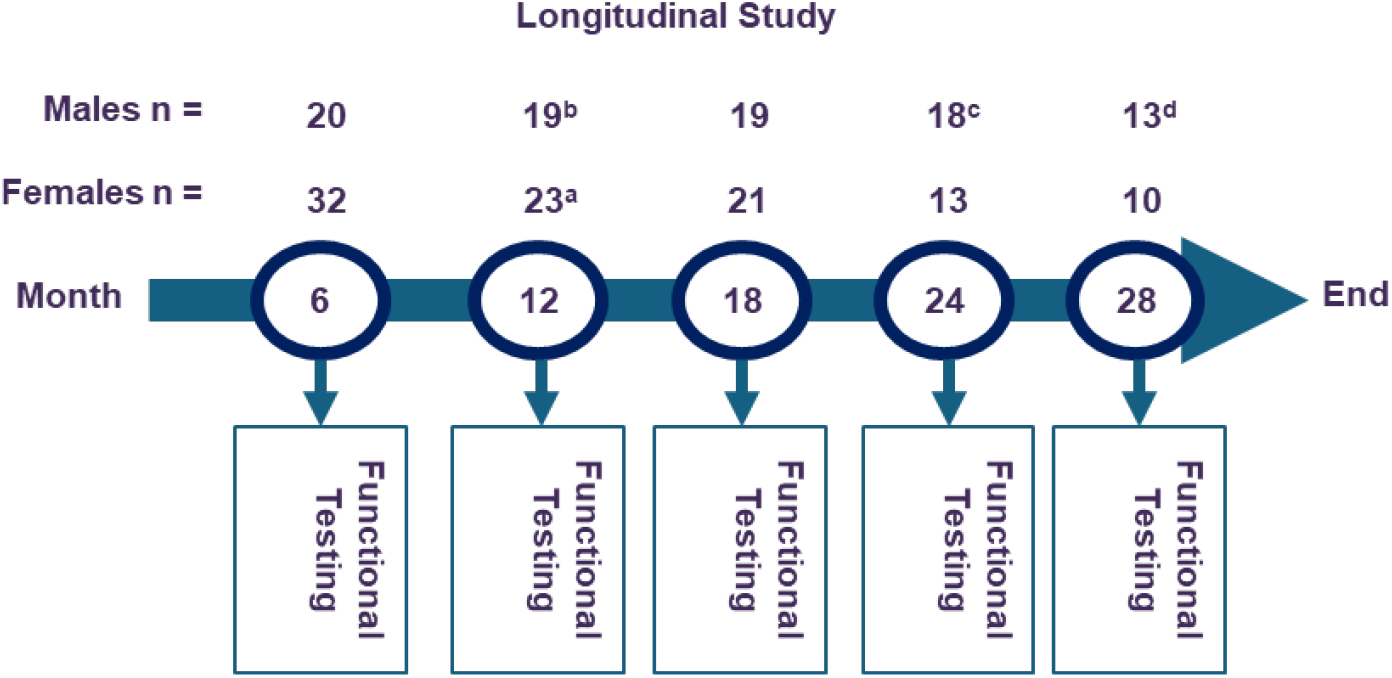
Study Design. Key: ^a^ 8 euthanized for tissue collection at 6m (months), ^b^ 1 euthanized for extreme dermatitis at 6m, ^c^ 1 dead of natural causes, ^d^ 5 dead of natural causes, functional testing includes CFAB (comprehensive functional assessment battery: rotarod, treadmill, inverted cling, voluntary wheel running, and grip test), collection of body weight, body composition (MRI for fat%) and in vivo contractile physiology (plantar flexor torque) in a subset of the mice.

### Physical Function

#### CFAB

Specific procedures for measuring physical function and calculating CFAB have been previously published.(10) In brief:

CFAB is a composite scoring system consisting of the sum of standardized determinant scores for each mouse based on mean and standard deviation for a 6-month-old (6m) sedentary control group, sexes calculated separately. A lower score indicates less functional aptitude in an individual mouse. A score of zero indicates a mouse performs as well as the average adult mouse, with a positive CFAB meaning the mouse has a higher function, and a negative score a lower function.

#### Previously Well-Validated CFAB Determinants

(10,13,14) See the Online Only Supplemental Methods section for a brief detailing of each CFAB determinant: rotarod (overall motor function), voluntary wheel running (volitional exercise rate and activity), inverted cling (overall strength/endurance), grip meter (fore-limb strength), and max speed treadmill test (aerobic capacity/endurance/speed).

#### Maximum Isometric Plantar Flexor Torque using *in vivo* Contractile Physiology

Details published elsewhere.(10,15,16) Briefly:

Mice were anesthetized with isoflurane to remove voluntary muscle control (VetEquip vaporizer), and then placed on heated platform of in vivo contractility apparatus (Aurora Scientific). Optimal subcutaneous electrode placement and current needed to produce maximal triceps surae plantar flexor twitch were determined. Maximal isometric torque (reported as mN/gbm) was then determined by applying that optimal current in 200 ms pulses in a 1 pulse, and 20, 40, 80, 100, 120, 150, and 180 Hz torque-frequency curve.

### Body Composition

#### Mass

The mice were weighed at each age point (scale = VWR-500P).

#### Fat %

Magnetic resonance imagery (EchoMRI) was used to measure lean and fat mass at each age of testing for females (6, 12, 18, 24, and 28 m) but males were only measured at 29m due to a combination of machine availability and the global pandemic lab shutdowns. Fat% We calculated fat percentage with the formula 100*((Lean mass + fat mass)/fat mass).

### Statistical Analysis

We used SPSS v29 (IBM) to analyze the data which is present as means ± standard error (se), unless otherwise noted (sd=standard deviation). Statistical significance is reported at p<0.05, trends at 0.05<p<0.10 and nonsignificant results (p>0.10) with ns.

We used repeated measures 1×5 ANOVA (5 time points) to evaluate change over time within the sexes, and report LSD (least significant difference) posthoc for a priori comparisons of 6m versus the older groups (Bonferroni adjustment also provided for comparison). We used a 2×5 mixed model ANOVA to compare the main effects of age and the sex*age interaction, with independent samples t-tests used to compare male to female means at each age for each variable. Effect size is reported as appropriate including partial η^2^ for ANOVAs and Cohen’s d for t-tests. Because of the great individual variability in CFAB determinant test scores at each age and the longitudinal nature of 5 age groups with 2 sexes, detecting differences in means is difficult (avoiding type 2 errors) when using strict multiple comparisons adjustments to avoid type 1 errors (i.e., Bonferroni). Thus, we provide both LSD and Bonferroni adjusted posthoc tests in the supplementary tables for the reader to evaluate. Certainly, prior work has established that in general age-related decline occurs in physical function so strict posthoc testing showing no difference should be critically evaluated. To compare the trajectories of decline, we curve fit a polynomial or linear curve. Nonparametric testing is used when appropriate (e.g., Friedman’s Two Way ANOVA by Ranks with Bonferroni posthoc). Statisical analysis including range, minimum and maximum values, mean, standard error, standard deviation, variance, skew, kurtosis, statistical tests for normality (Shapiro-Wilks and Kolmogorov-Smirnov), and parametric means testing and/or nonparametric tests (Friedman’s 2-Way ANOVA by Rank), as appropriate, for each of the variables below are presented in **Supplementary Tables S1 Males, S2 Females, and S3 Males versus Females**.

## Results

### CFAB

We used our CFAB composite scoring system to determine overall physical function and exercise capacity at each age that we evaluated in both males and females. A negative CFAB score indicates worse function compared to the 6m mean.

- Male CFAB mean reduced -1.6 from 6m to 12m (trend, p=0.091), -3.6 from 6m to 18m (p=0.001), -5.9 from 6m to 24m (p<0.001), -7.1 from 6m to 29m (p<0.001), -2.0 from 12m to 18m (p=0.041), -2.3 from 18m to 25m (p=0.006), and -1.2 from 25m to 29m (p=0.005). *Test:* 1×5 repeated measures ANOVA, F = 26.9, p<0.001, partial η^2^ = 0.69, LSD posthoc. For more details see **Figure 2A**, and **Table S1**.
- Females CFAB mean declined by -10.6 from 6m to 12m, -15.2 from 6m to 18m, -20.4 from 6m to 24m, and -20.8 from 6m to 28m; all at p<0.001. The rate of decline plateaued at the older ages, however: -4.5 from 12m to 18m (p<0.001), -5.21 from 18m to 24m (p<0.001), and only -0.61 from 25m to 28m (ns). *Test:* 1×5 repeated measures ANOVA, F=26.9, p<0.001, partial η^2^=0.69, LSD posthoc. For more details see **Figure 2C**, and **Table S2**
- The was a linear association overall between the data in both males and females. The slope of functional decline in females was more than 3x steeper (slope = -1.0 CFAB/month, R=0.88) than that of males (slope = -0.31 CFAB/month, R=0.71). For more details see **Figure 2B and 2D**. Furthermore, by examining the overall change in means longitudinally, we found that in males there was a steady linear decline of function (regression equation: CFAB= -0.315months+2.025, R=0.999), whereas the female mean regression fit better as a 2^nd^ degree polynomial (regression equation: CFAB= 0.0379months^2^ – 2.223m + 11.613, R=0.993) than linear (R=0.93). The slope of the male decline was quite steady (∼0.32), but the decline was most accelerated for females between 6 and 12m (slope=-1.77), reduced from 12 to 24m (slope=-0.79), and stalled between 24 and 28m (slope=-0.22). See also **Figure S1 A and B** in the Supplemental Section.

**Figure 2.**
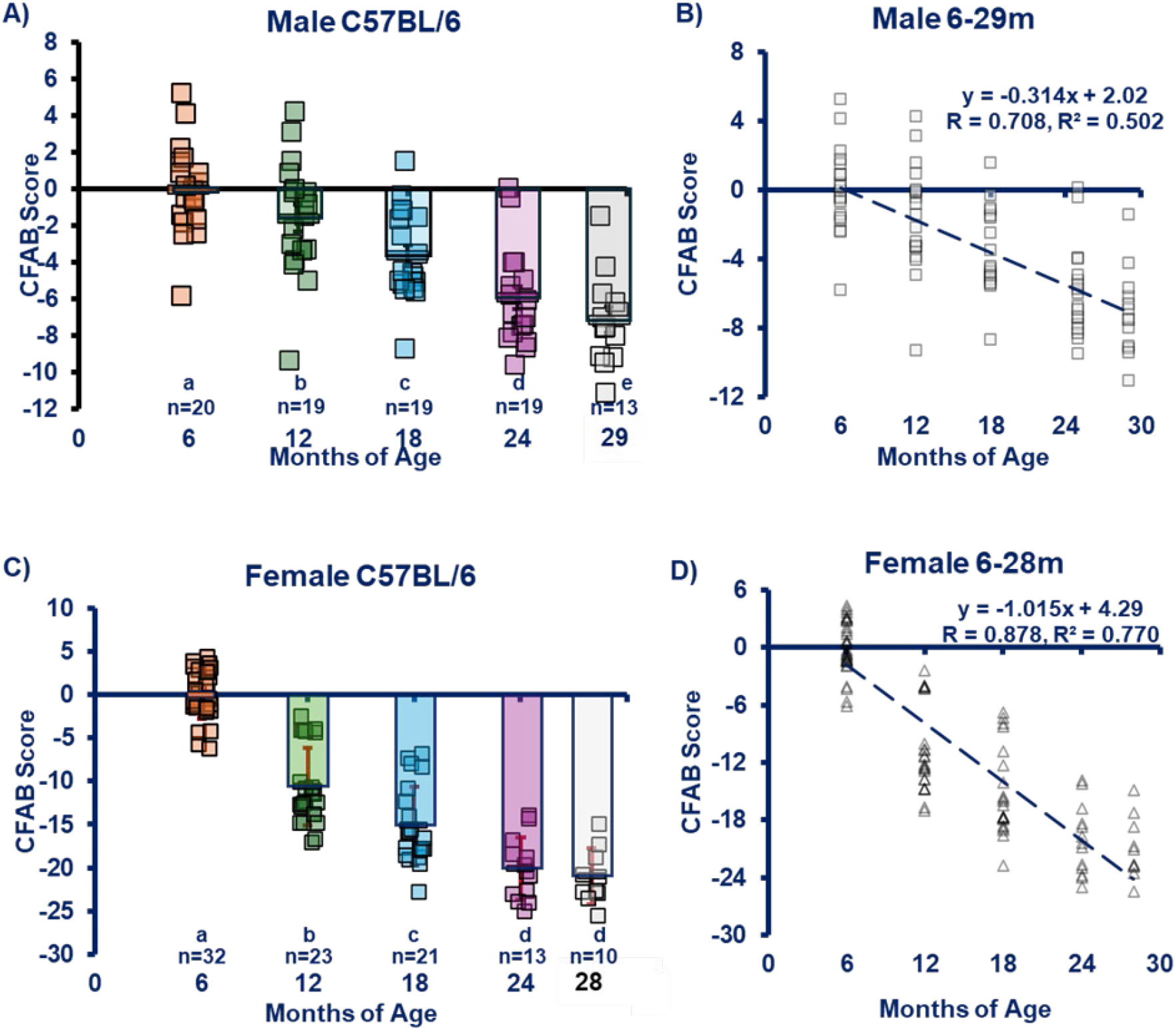
CFAB (Comprehensive Functional Assessment Battery). CFAB is a composite scoring system measuring overall exercise capacity and physical function. The average adult would have a score of zero, score less than zero indicate worse function in compared to the average adult. **A) Males. B) Females. C) Male Linear Regression**. Males follow a very linear decline with age (−0.3 CFAB per month). **D) Female Linear regression**. Females show an accelerated slope of decline (−1 CFAB per month, 3.33x compared to males). Key: m = months of age, Equations and regression lines = linear regressions. Each symbol (triangles in Panel A and circles in Panel B and C = the mean CFAB score of all the surviving mice at the given age). Different letters = significant difference between groups, same letter = no significant difference

### CFAB Determinants

VWR (Activity rate and volitional exercise, **Figure S2**), rotarod (overall motor function--coordination, balance, endurance, and power, **Figure S3**), grip test (forelimb strength, **Figure S4**), inverted cling (four-Limb strength/endurance, **Figure S5**), and treadmill max speed test (aerobic capacity and endurance, **Figure S6**) are the tests that make up CFAB. In brief:

- Male means *numerically* declined, though not significantly (ns) at a similar rate (−9-10%) from 6 to 12m in normalized grip, rotarod, treadmill, and cling. However, VWR mean was -34% (ns), and cling*gbm did not change (ns, -1%). After 12m, most of the determinants significantly declined more rapidly Treadmill however did not decline significantly compared to 6m until 25m (−42%) and 29m (−38%), p<0.001 in both cases. VWR had the largest changes of -62%, -75%, and -89% at 18, 25 and 29m, respectively. See **Tables 1 and S1** for details.
- Females means declined from 6 to 12m in normalized grip (−34%, p<0.001), rotarod (−29%, p=0.011), treadmill (−11%, p=0.078), cling (−48%, p<0.001), VWR (−62%, p=0.07) and Cling*gbm (−30%, p=0.003). As the females got older there were significant decreases from the 6m mean with all other age groups other than treadmill, which similarly to the males did not decrease (−9%, p=0.388) at 18m. See **Tables 1 and S2** for more details.
- Males were remarkably similar to females in treadmill both in means and in percent decline with age over the lifespan (no significant differences). However, there were some large sexual dimorphisms in the other determinants. See **Table S3** for more details.
  ∘ For example, in VWR (m) means at 6m (m=1639±1053sd, f=4730±3359sd) were 2.9 fold greater in females (p<0.001). As the mice aged however, male VWR performance began to catch up (female means were only 1.67 and 1.4 fold greater at 12 and 18m, p=0.078 and ns respectively). At 25 and 29 months, males (m) outperformed female (f) mice ∼2 fold at both 25m (mean meters traveled per day: m=410m±330sd, 24m f=197±339sd, p=0.086) and 29m (means: m=172m±170sd, 28m f=85±3124sd, p=0.186).
  ∘ Females were much stronger than males in normalized grip (mN/gbm) at 6m (m=44.9±9.0sd, f=62.2±9.0sd, p<0.001), though the differences equaled out at more advanced ages (no significant differences). However, in rotarod males and females were similar at 6m, but both declined at later ages with males maintaining higher overall function (fold change 1.4, 1.3, 1.6, and 1.9 with pval = <0.001, 0.094, 0.039, and 0.036 at 12, 18, 25/24, and 29/28m respectively).
  ∘ Finally, inverted cling*gbm demonstrated that females (10468±1507) began with much greater ability (p<0.001) at adulthood than males (4584±2364). Importantly, the basic inverted cling test has a ceiling of 420 seconds. No males reached the ceiling but most 6m females did. The females (−30%, p=0.004), however, declined more rapidly in cling*gbm performance at 12m than males (−1%, ns), even with female 6m performance capped at the ceiling. In the 12m females only four reached the ceiling measurement out of 23 tested. Males and females were not significantly different at 12 and 18 months. Interestingly, male mice had significantly higher scores at the oldest two ages than females (p=0.030 and 0.028, respectively). This suggests that male strength was better preserved during aging even though females started out stronger.

**Table 1.**
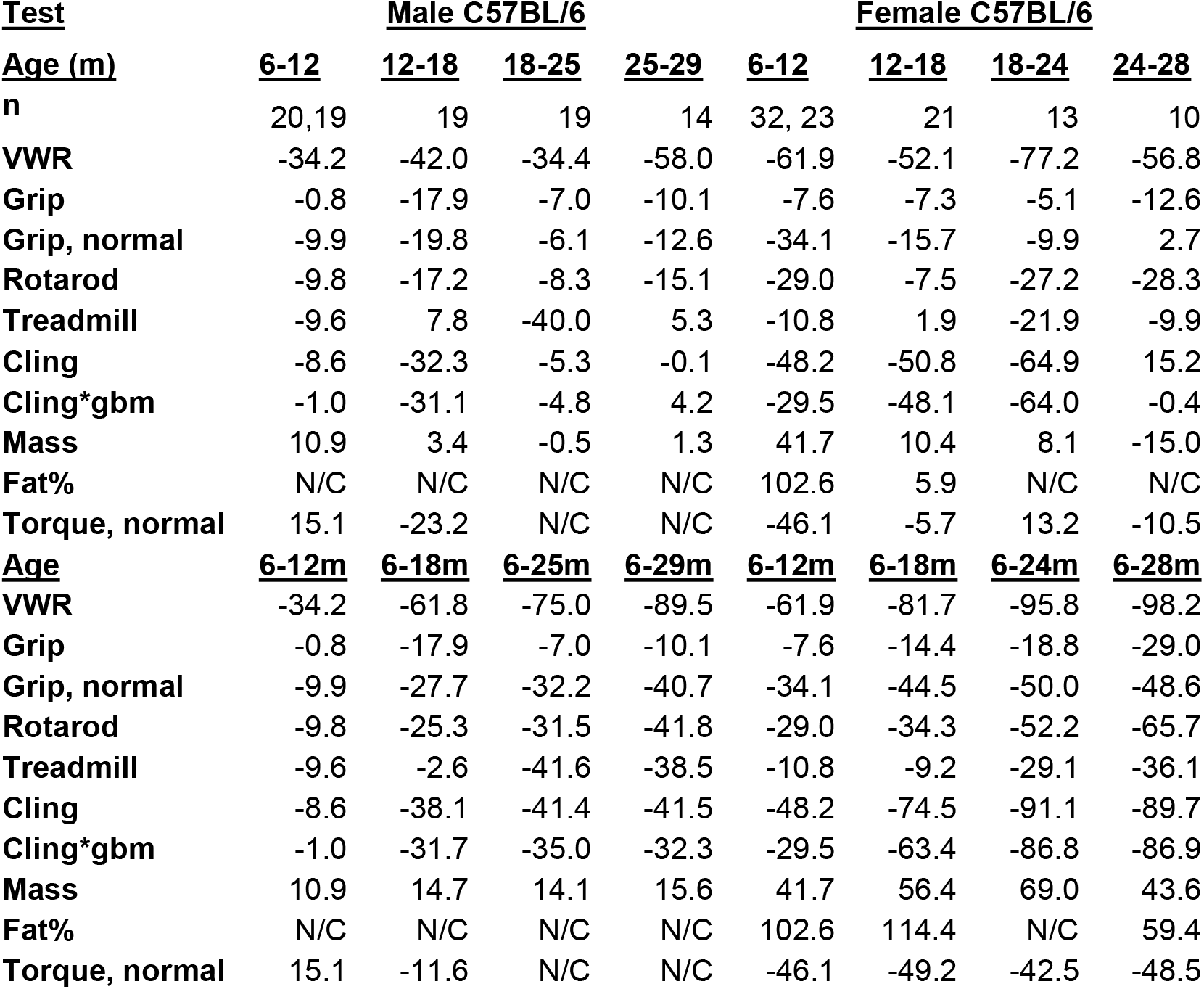
Percent Change of Mean. Males are in the first set of data columns (2-5) and females are in the second set (6-9). **Key:** VWR = voluntary wheel running, gbm = grams body mass, normal (normalized) = divided by gbm, * = multiplied by, m = months, each number = the numerical percent change in means from the first listed month to the second, N/C = not collected, n = number of mice in age group with 6-12m listing first 6m then 12m, and the rest listing only the second month.

### Contractile Physiology

We assessed a subset of mice for maximum isometric plantar flexor torque as a measure of muscle contractility. The groups were not balanced and some data points were missed (most 25m and 29m males), due to restrictions during the global pandemic. We report the results normalized to body mass with statistics from repeated measures ANOVA with LSD posthoc for males and females. The comparison of males to females used independent samples t-tests. See **Table 1, Figure 3**, and **Tables S1-S3** for details.

**Figure 3.**
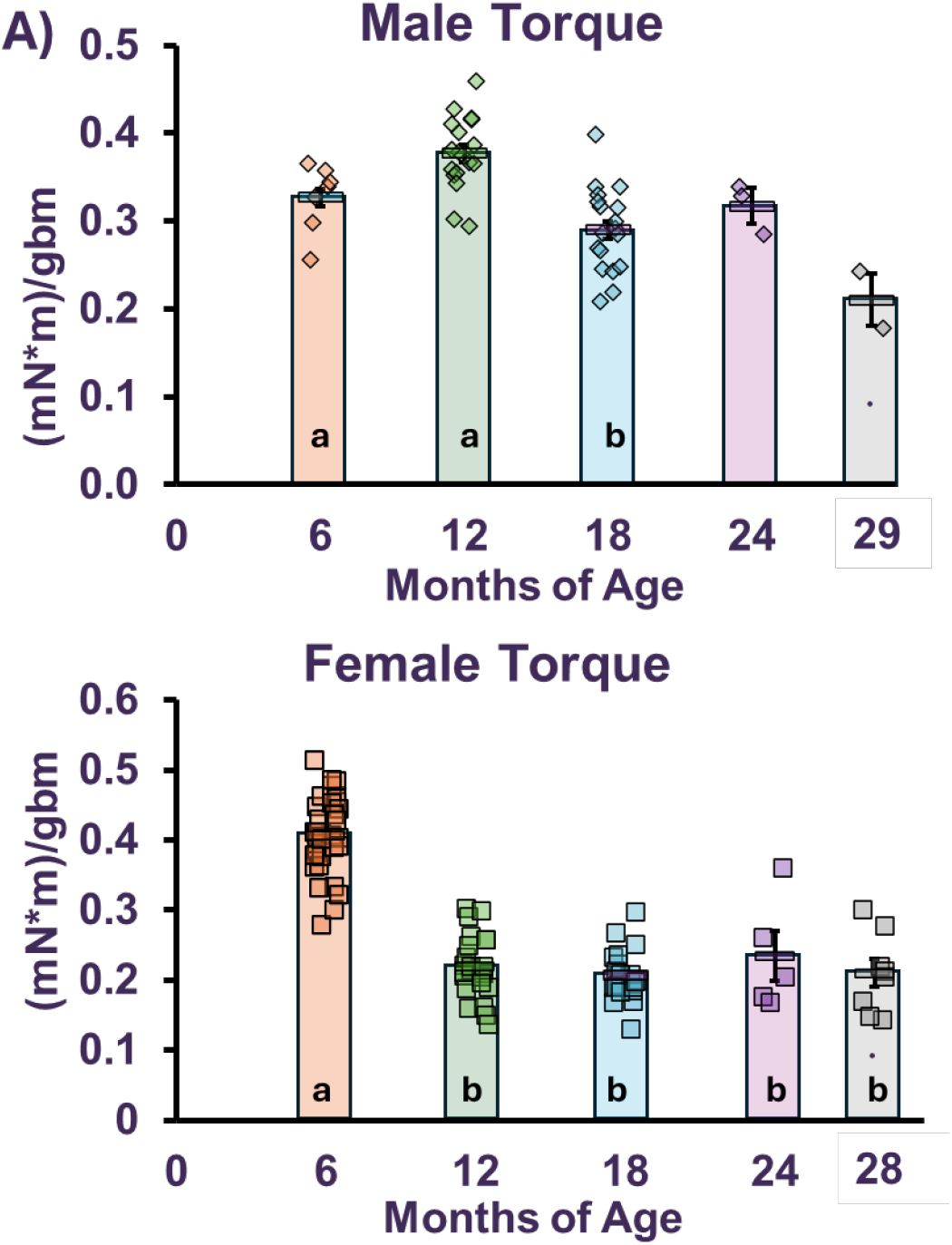
Plantar Flexor Torque. Normalized to GBM (grams body mass). A) Males. Stats only 6-18m. B) Females. **KEY:** Each symbol = 1 mouse. Different letters = statistically significant (repeated measures ANOVA) p<0.05

- In males mean torque did not alter significantly from 6-12m, but then reduced from 12-18m (p=0.035).
- Female mean torque reduced at 12m (p<0.001) and then remained unchanged at the older ages.
- Comparing mean male torque to female torque at 6-18m ages (ages where more complete data sets are available for each sex), demonstrated that males were weaker than females at 6m (percent difference 21), but then were stronger at 12 (52%) and 18m (33%), all comparisons with p<0.001.

### Body Composition

All surviving mice were weighed at each time point and females (as well as 29m old males) were evaluated for body composition with MRI. See **Table 1**. We report the results normalized to body mass with statistics from repeated measures ANOVA with LSD posthoc for males and females. The comparison of males to females used independent samples t-tests. **Figure 4** and **Tables S1-S3** for details.

**Figure 4.**
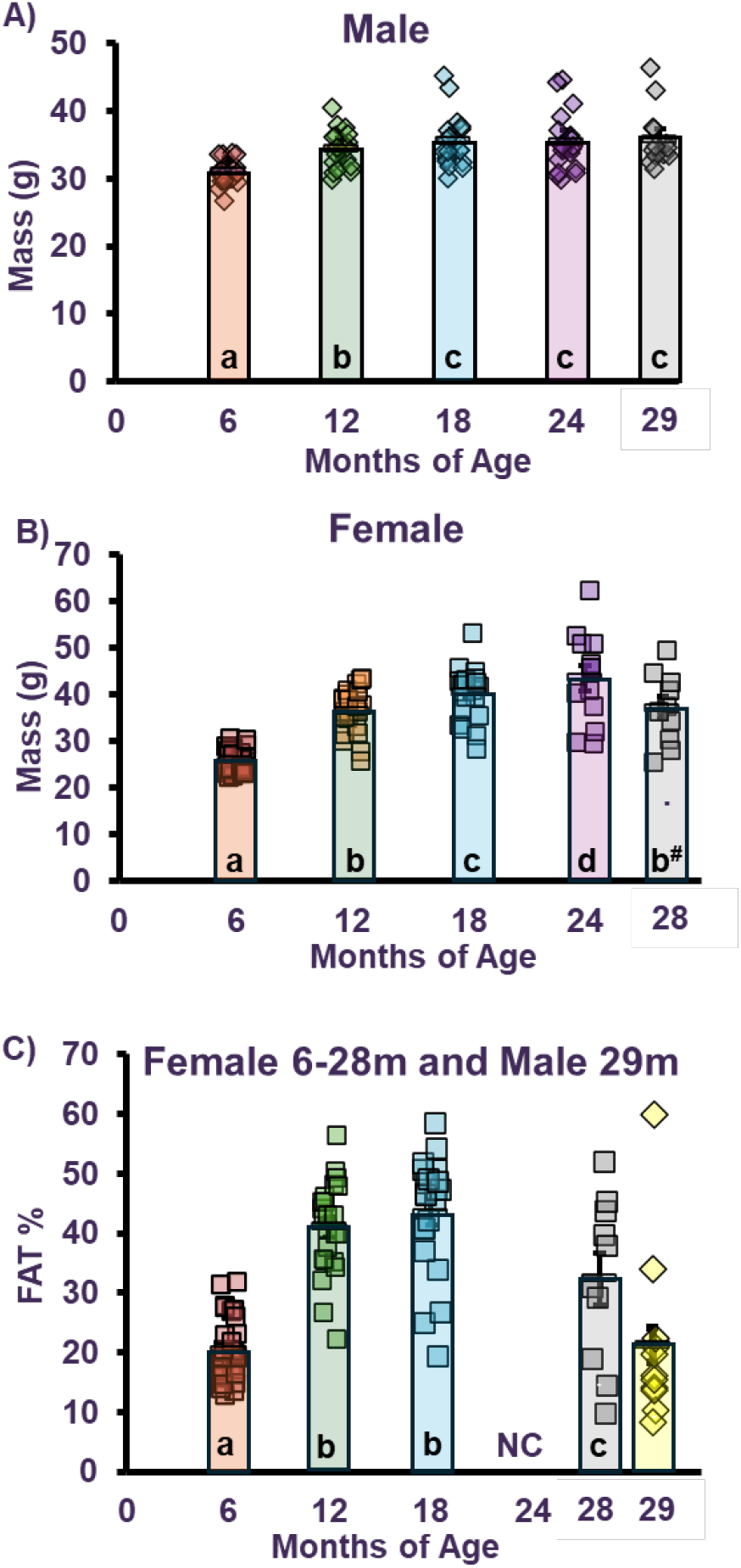
Body Composition. A) Males Body Mass. B) Females Body Mass. C) Fat Percentage. All mice are female, except @ 29m (male in diamonds). KEY: Each symbol = 1 mouse, error bars = standard error, NC = not collected (covid restrictions). Different letters indicate statistical significance at p<0.05.

*Mass:*

- Male mice gained weight from 6-12m and from 12-18m (p<.001 and p=0.004, respectively, but then remained the same weight up to 29m (ns: 18-25m and 25-29m).
- In contrast, the female mice gained weight from 6 to 12m (+42%, p<0.001), then steadily gained up to 24m (12-18m +10%, p<0.001; from 18-24m +8%, p=0.011), before losing 15% of their weight from 24-28m (p<0.001).
- Comparing the male to female mice mass revealed significant differences in patterns of weight gain. Females gained much more weight overall from 6-24m than males from 6-25m (male +4.4g vs. female +18.8g), even with males starting out heavier (+18% difference, p<0.001). At the oldest age males and females had similar mass (35.8±1.0g for males and 37.0±2.4g for females, ns), due to surviving females losing weight.

*Fat Percentage:*

- Due to events beyond our control only the 29m male mice were assessed for fat percentage.
- Female mice increased fat% by 21 percentage points from 6 to 12m (p<0.001), then remained the same from 12-18m (ns). We were unable to measure the 24m timepoint for the females, but from 18-28m the surviving mice reduced fat% by 26% (p=0.016).
- Mean male fat% at the oldest ages tended (p=0.095) to be lower than female (21.1±4.4% for males and 32.3±4.5% for females, Cohen’s d=-0.079).

## Discussion

### Physical Performance

Overall physical function measured by CFAB declined with age in both males and females. As hypothesized there were differences in the rates of decline and the decline patterns of individual parameters (e.g., rotarod, grip, cling, mass, etc.) between the two sexes.

Previously we have used cross-sectional studies to demonstrate age-related functional decline in male mice. The neuromuscular healthspan scoring system (NMHSS; consisting of: rotarod, inverted cling, and *in vitro* EDL maximum isometric force) was a composite scoring system that accounted for the vast individual variability in scoring of functional tests by utilizing multilinear regression equations to build a term that improved power of detection markedly compared to the individual determinants.(13) We followed up by reverse translating the Fried Frailty Phenotype showing that mouse frailty occurs at a similar rate and life stage to humans.(17) Then, during a study in which we demonstrated wheel running could reverse frailty, we adapted our frailty phenotype into the precursor of CFAB, the intervention assessment value (IAV), to detect pre-to post-exercise changes in frailty measurement.(14) Finally, we established and validated CFAB in male C57BL/6 mice, and then used it to measure exercise intervention efficacy.(10,12) This lead to the current study where we expanded CFAB to females and lifespan longitudinal design.

We know that in general overall physical function in mice declines with age, though specifics vary between researchers, studies, measurements, and strains.(18) For example, Justice and colleagues showed that overall composite motor function declined 7.4% from 3 to 20 months and then declined accelerated by reducing a further 13.5% from 20 to 26 months.(19) Gait speed decline over the lifespan was shown to be fully explained by frailty status and age in a study by Petr and colleagues.(20) A recent preprint by Liao and colleagues(21) compared C57BL/6 and CB6F1 (outbred hybrid of C57BL/6 and Balb/c), noting that age-related changes to physical and cognitive function had altered trajectories depending upon strain. They found that in C57BL/6 there was no change between 4 and 12m in rotarod, grip or VWR, but in CB6F rotarod and grip declined during the same period (VWR did not).(21) Previously we determined that C57BL/6 males had overall declining function (measured by CFAB) between 6 and 10m and between 24 and 28m, with different determinants changing at different rates in the two age groups (e.g., rotarod declined by 37% in the adults but only by 10% in the older adult group).(12)

While traditionally many studies of physical function during aging have been done in male C57BL/6, there have been numerous studies done in other strains and in females, and some longitudinal.(22–25) The SLAM (study of longitudinal aging in mice) study at NIA uses both C57BL/6J and the HET3 outbred strain, and notes that rotarod longitudinal performance is enhanced in longitudinal versus cross-sectional studies, presumably by a training/learning effect.(26) Some studies show that certain parameters (i.e., rotarod) do not change with age, which can sometimes be attributed to sexual dimorphisms. Fisher, et al. 2016 found females did not have longitudinal changes in rotarod from 4m to 20, 24 or 28m but males did.(23) However, in contrast, Kwak and colleagues showed females in a longitudinal frailty study having lower rotarod scores at 23 and 26m compared to 17 and 20m.(27) Our current study found that female rotarod performance was reduced from 6m to 12m remained static at 18m, reduced again at 24m and didn’t change at 28m; but males declined the most from 12 to 18m and then from 25 to 29m.

The effects of aging on physical performance in mice are in the context of strain, sex, and study characteristics. By establishing longitudinal baseline results using C57BL/6 male and female mice, and our CFAB system, in future efforts we can recognize and identify causal factors of reduced physical function.

### Contractile Physiology

Numerous cross-sectional studies (typically measuring the EDL, extensor digitorum longus, and SOL, soleus) have reported age-associated declines in whole muscle force, power, and velocity of contraction.(28–32) Since *ex vivo/in vitro* whole muscle contractile physiology is by necessity a terminal procedure, *in vivo* contractile physiology as a repeatable measure is an alternative. There has been some work using torque-production as a measure of function in short-term intervention or cross-sectional aging studies.(10,15,33,34) For instance, Sheth and colleagues showed both males and females having lowered normalized plantar flexor torque at 20m versus 8 or 12m.(35) However, beyond the current study, there has been little longitudinal investigation of age-related changes of *in vivo* contractile physiology (e.g., plantar flexor or dorsiflexor torque) in mice and this remains a much needed area of further investigation. Importantly in our study we note that female torque output adjusted for body mass remains remarkably similar over the lifespan after a large initial drop at 12m. More work is needed to understand this observation in the context of sarcopenia, as it may be a consequence of survivor bias not observed in cross-sectional studies.. Future mechanistic work will connect CFAB, muscle contraction, and aging.

### Body Composition

Patterns from the literature of age-related changes to body mass and fat% differ based on sex and often whether a study is cross-sectional or longitudinal. Our current work demonstrated sexual dimorphism in weight gain/maintenance with females steadily gaining weight up to 24m then losing weight at 28m, while male mass increased up to 18m and then remained static from 18-29m. In females we noted a substantial increase in fat deposition from 6 to 12m, but later on there was no change at 18m with a reduction in fat in the surviving mice at 28m. It has been previously reported that female mice gain fat before losing it at older ages.(27)

Interestingly, there were strong correlations (R=-0.79, p<0.001) between body mass and CFAB and body mass with age (R=0.76 from 6m to 24m, p<0.001) in females but a more moderate relationship with mass and CFBA in males (mass vs. CFAB, R=-0.622, p<0.001). There was minimal relationship between age and mass in males (R=0.405, p<0.001). See **Figure S7** in the supplement for more details. This might account for some of the more rapid decline of CFAB observed in females as they gained large amount of weight that was mostly fat up through 24m.

### Female Aging

One reason that 12m may be such an important inflexion point for female mice is that by around 9-12m of age mice begin transition to the so-called estropause where they experience irregular estrous cycles which cause hormonal fluctuations similar to human menopause with a dysregulation in the hypothalamic–pituitary–gonadal endocrine axis.(36) Estrogen has been found to play a critical role in female mouse strength maintenance and declining levels.(37) Whether via ovariectomy, or natural aging, declining estradiol levels induce proteomic changes related to muscle contraction and structural integrity.(37,38) In addition, there is a well-known association between menopause and weight/fat% gain in middle-age humans which may be contributing to the sexual dimorphism in body mass change in the current study.(39,40) However, mouse models do not perfectly model human menopause because at some point during advanced aging ovarian function is spontaneously restored in roughly 75% of mice, and this spontaneous restoration of hormonal homeostasis may account for some of the improvements in function and body composition seen at the oldest ages in our study.(41)

### Caveats

This project was influenced by the SARS-CoV-2 worldwide pandemic. Due to lab shutdowns some data points (contractile physiology on older males for example) were unable to be collected on some of the mice. Additionally, when this work was started, with the male cohort, we did not have access to devices to measure fat% (e.g., EchoMRI), thus those data points are missing.

Males and female had quite different scales for CFAB and so we did not compare them directly. However, we did compare the rates of decline (slopes of regressions) and the individual determinants, demonstrating that there was sexual dimorphism in functional aging patterns. Females also died at a much higher rate than males with

## Conclusion

Both males and females steadily lost physical function during aging. However, there were sexual dimorphisms in the patterns of functional loss and in the rates of change for many of the parameters as well as CFAB overall. We accomplished primary purpose to establish longitudinal baselines and validate CFAB in female mice. Going forward, more research is needed to establish the mechanisms that account for differential function aging. At each age group the marked differences in CFAB scores for the upper quintile, which are similar in performance to the younger age mean, and the lower quintile demonstrate that those mice in the higher performing groups may be experiencing so-called “successful aging.” We previously examined transcriptome differences associated with CFAB between different age-groups(11), but a multiomics comparison of high and low functioning individuals within age cohorts is a natural future step to hunt for potential novel mechanisms. In conclusion, CFAB has been shown to be a valuable aid for measuring overall physical function and exercise capacity in males and females and in longitudinal, cross-sectional, and intervention studies.

## Supporting information

Supplemental Section

## Animal Use Statement

All animals used in this study were treated humanely under East Carolina University IACUC guidelines and protocol #p106a.

## Author Contributions

The authors recognize the relative contributions as: Conceptualization, TGG; Methodology, TGG; Validation, TGG, RM; Formal Analysis, TGG, RM, MLP; Investigation TGG, MLP, RM, CB; Resources TGG; Writing – Original Draft TGG; Writing – Review & Editing TGG, RM, MLP, CB; Supervision TGG, MLP; Project Administration TGG; Funding Acquisition TGG.

## Acknowledgements

Chris Byrd, DPT is now a practicing clinician. Megan L. Pajski, PhD is now at ThermoScientific.

## Online Only Supplemental Section for

**<u>Table of Contents</u>**

**Page Topic**

**S1 Supplemental Section Methods**

**S2 Figure S1 CFAB Mean Regresssions**

**S3 Figure S2 VWR**

**S4 Figure S3 Rotarod**

**S5 Figure S4 Grip Test**

**S6 Figure S5 Inverted Cling**

**S7 Figure S6 Treadmill Max Speed Test**

**S8 Figure S7 Body Mass Regressions**

**S9 Supplemental Section References**

**Excel File S1 Table S1 Male Statistics and Descriptives**

**Excel File S2 Table S2 Female Statistics and Descriptives**

**Excel File S3 Table S3 Male versus Female Statistics and Descriptives**

