## Supplemental Section for "Longitudinal Decline of Exercise Capacity in Male and Female Mice"


### Table of Contents


| Page | Topic |
|---|---|
| S1 | Supplemental Section Methods |
| S2 | Figure S1 CFAB Mean Regresssions |
| S3 | Figure S2 VWR |
| S4 | Figure S3 Rotarod |
| S5 | Figure S4 Grip Test |
| S6 | Figure S5 Inverted Cling |
| S7 | Figure S6 Treadmill Max Speed Test |
| S8 | Figure S7 Body Mass Regressions |
| S9 | Supplemental Section References |
| Excel File S1 | Table S1 Male Statistics and Descriptives |
| Excel File S2 | Table S2 Female Statistics and Descriptives |
| Excel File S3 | Table S3 Male versus Female Statistics and Descriptives |

**Online Only Supplementary Methods Section:**

**Previously Well-Validated CFAB Determinants:** CFAB determinants include the following: rotarod (overall motor function), voluntary wheel running (volitional exercise rate and activity), inverted cling (overall strength/endurance), grip meter (fore-limb strength), and max speed treadmill test (aerobic capacity/endurance/speed). Details have been previously published.(1–6) Briefly:

- Voluntary wheel running (volitional exercise and activity rates): Average meter/day over 1-week reported (Columbus Instruments)
- Rotarod (overall motor function: balance, stamina, coordination, power): After 2 acclimation sessions, best of 3 trials latency to fall (seconds, sec) while accelerating @ 4-40 rpm/5 minutes (PanLab).
- Grip Meter (forelimb strength): best of 5 trials recorded in Newtons (N), reported as N/gbm (grams body mass) (Bioseb)
- Treadmill (Aerobic capacity): After two sessions of acclimation, on day 3 the outcome is time to failure (sec) from an initial velocity of 4 m/min accelerating at 0.6 m/ min / 20 seconds. (Columbus Instruments)
- Inverted Cling (overall strength / endurance): Best of two trials, latency to fall is the outcome measurement (sec). (Custom built device)

For females the VWR and cling measurements violated normality severely and thus we used a log10 transformation of VWR (meters run per day) and normalized cling (cling time in seconds * grams body mass) to standardize for the CFAB measurement. Rotarod, treadmill, and grip had normal distributions. See also **Supplementary Table S2.**

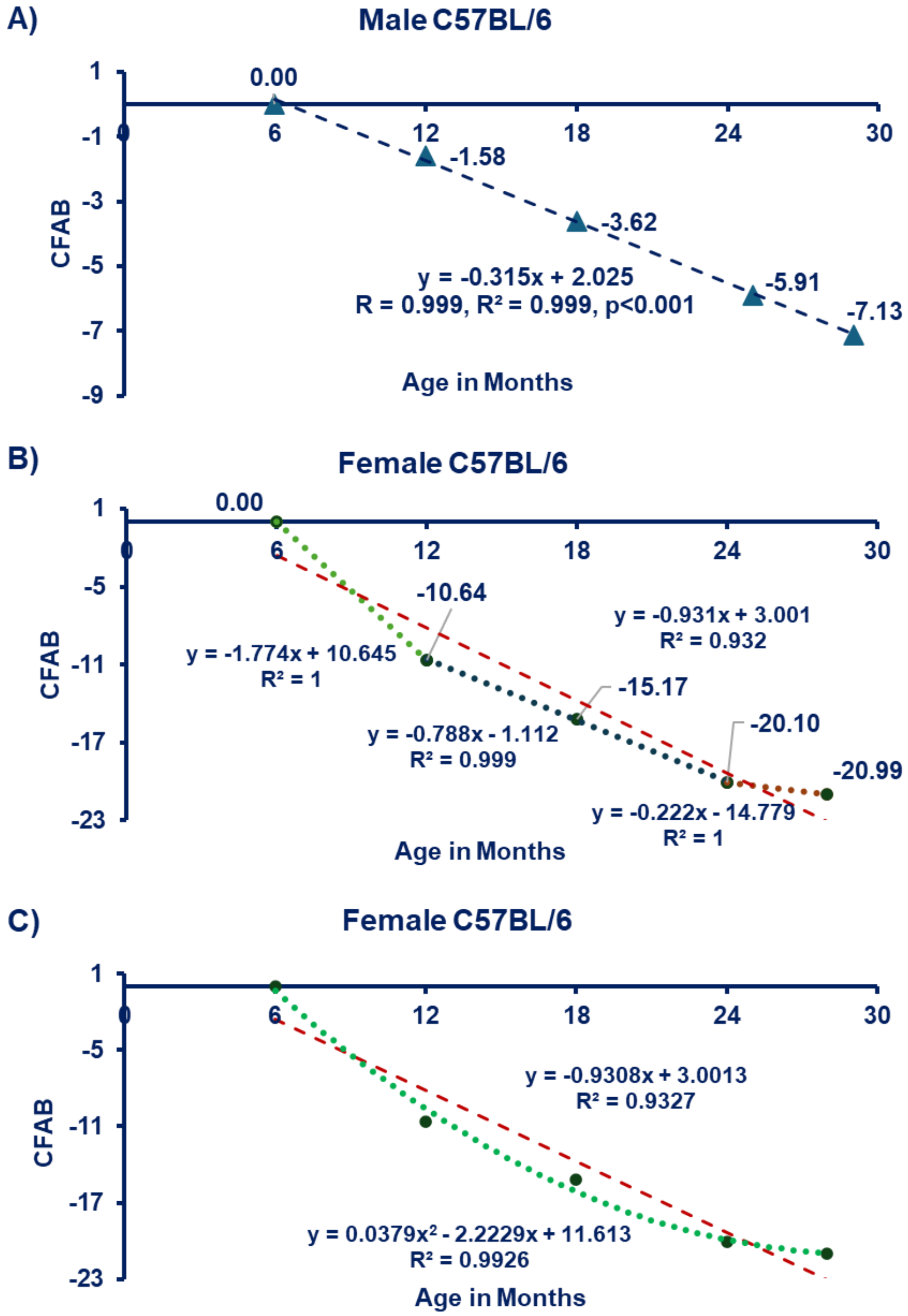


**Figure S1 CFAB Mean Regressions with Age. A) Males Linear.** Males follow a very linear decline with age (-0.3 CFAB per month). **B) Females Linear.** Females show an accelerated slope of decline compared to males that alters with age, with a larger decline from 6 to 12m (-1.77 per month, roughly 5.6 time the rate of decline of males), that slows down between 12 and 25m (-0.788 per month, 2.5x the rate of males), and then slows even more between 24 and 28 months (-0.222 per month, about 70% slower than the overall male rate. C) Female Linear and Polynomial. The linear fit is inferior to the polynomial fit, given the changing rate of decline between different ages. Key: m = months of age, Equations and regression lines = best fit regressions (in Panel A and B = linear; Panel C shows both best linear and polynomial regression), Each symbol (triangles in Panel A and circles in Panel B and C = the mean CFAB score of all the surviving mice at the given age).

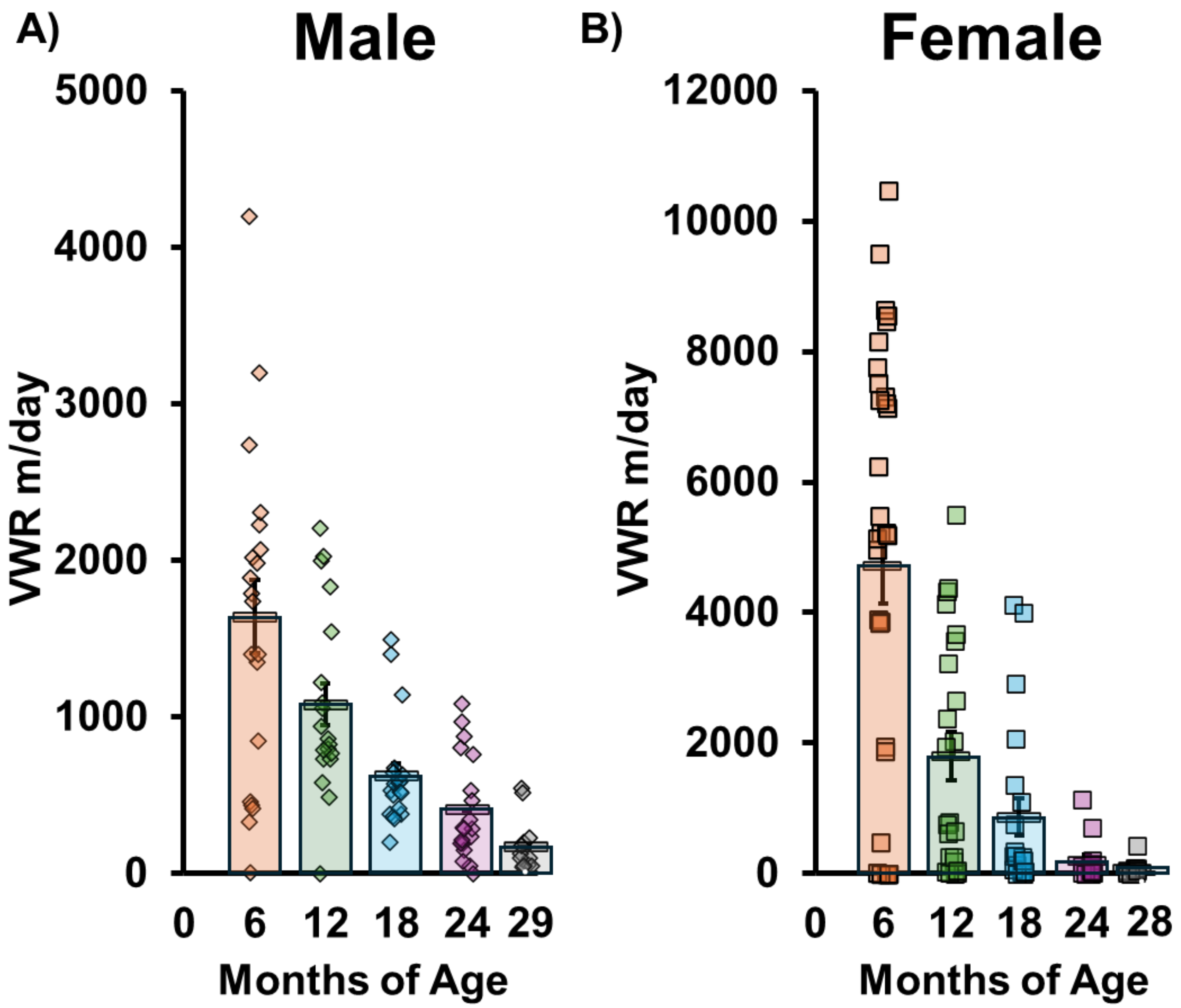


**Figure S2 VWR (Voluntary Wheel Running). A) Males.) Females. Key:** m = meter, Each symbol (diamonds in Panel A and squares in Panel B) = the VWR distance ran per day of all the surviving mice at the given age. Different letters = significant difference (i.e., same letter = no difference in means), # = , error bars = standard error, columns = mean.

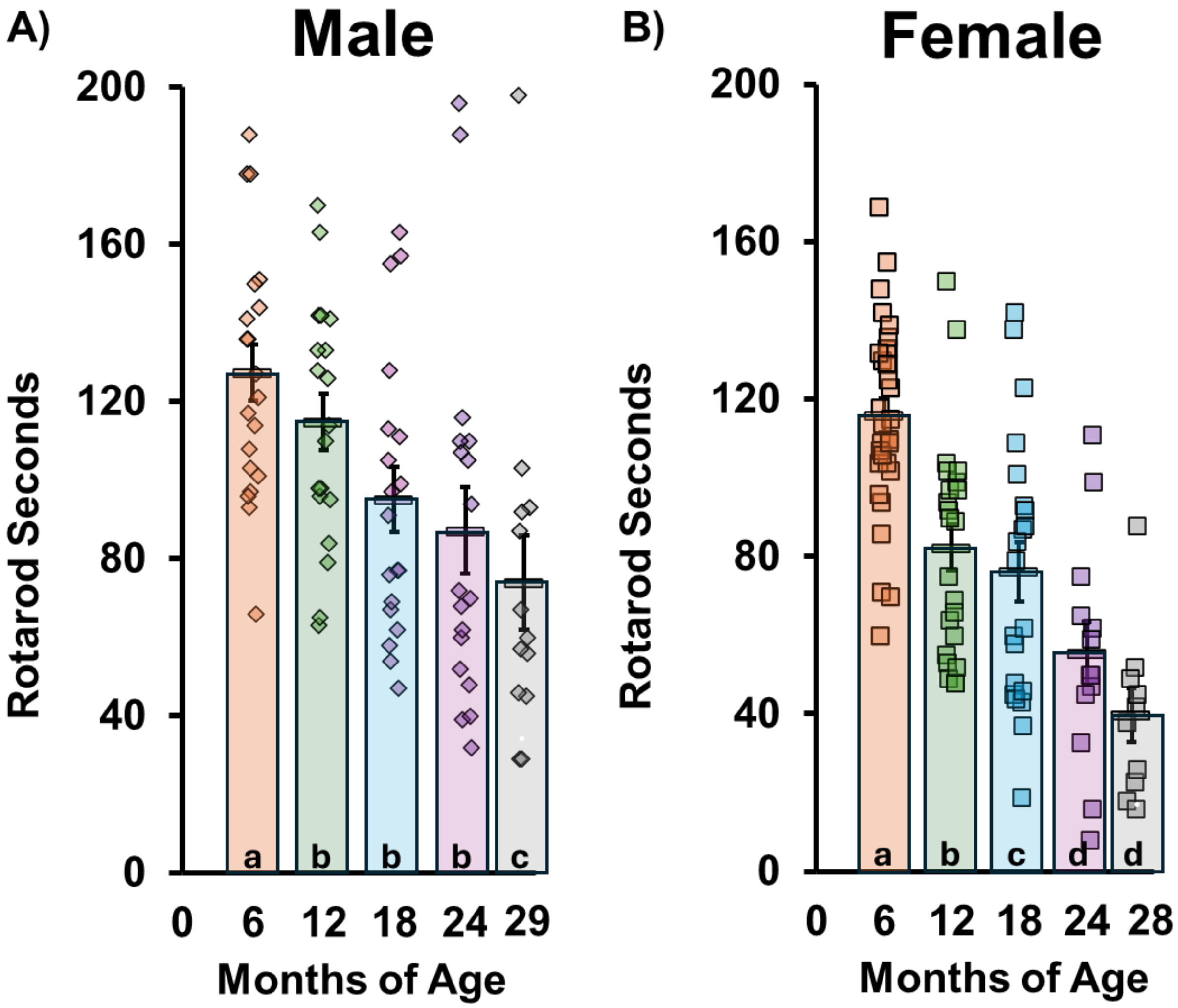


**Figure S3 Rotarod. A) Males B) Females. Key:** Each symbol (diamonds in Panel A and squares in Panel B) = the rotarod latency to fall of all the surviving mice at the given age. Different letters = significant difference (i.e., same letter = no difference in means), error bars = standard error, columns = mean.

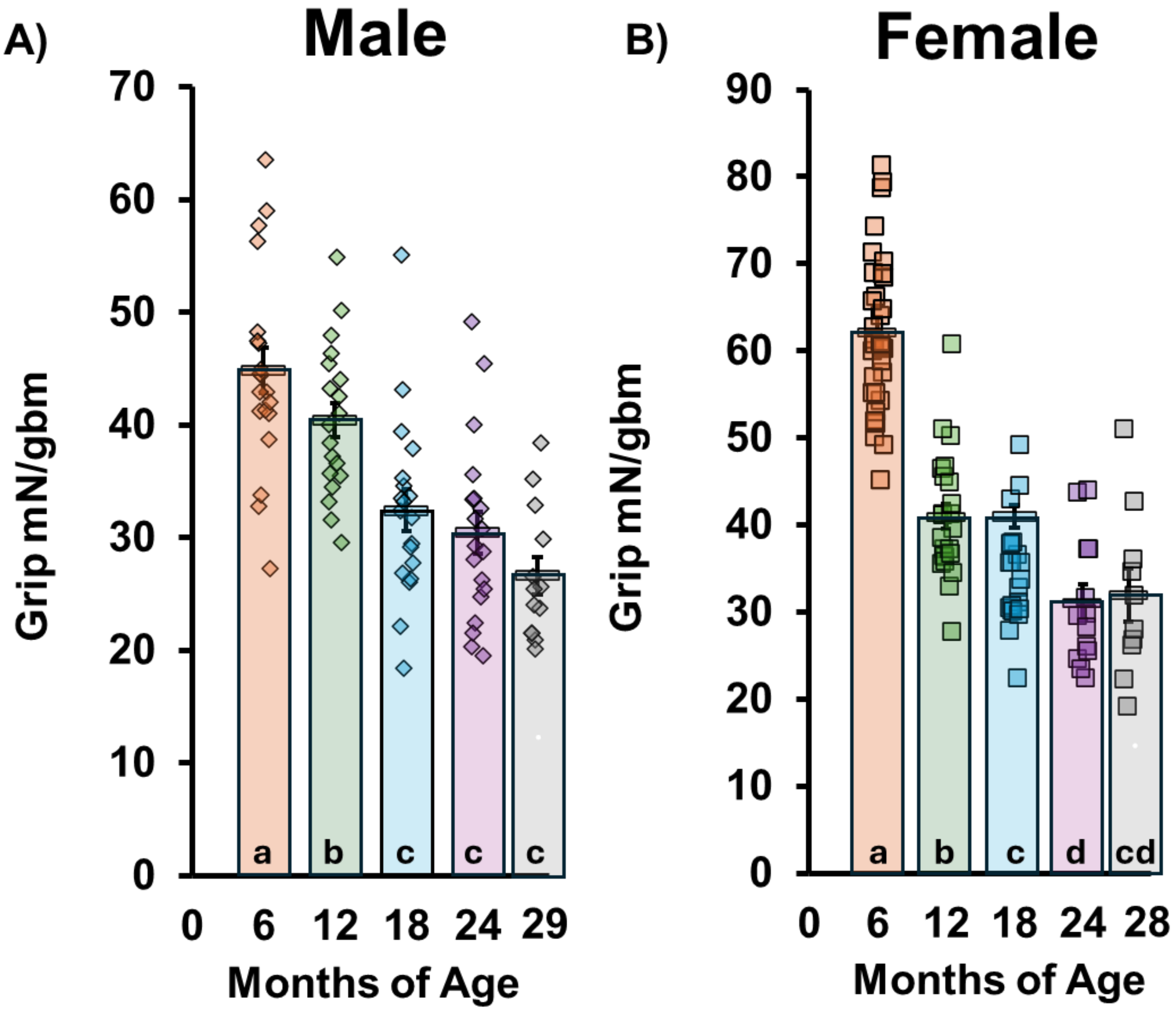


**Figure S4 Grip Test. A) Males B) Females. Key:** mN = milliNewton, Each symbol (diamonds in Panel A and squares in Panel B) = the grip strength normalized to grams body mass (gbm) of all the surviving mice at the given age. Different letters = significant difference (i.e., same letter = no difference in means), error bars = standard error, columns = mean.

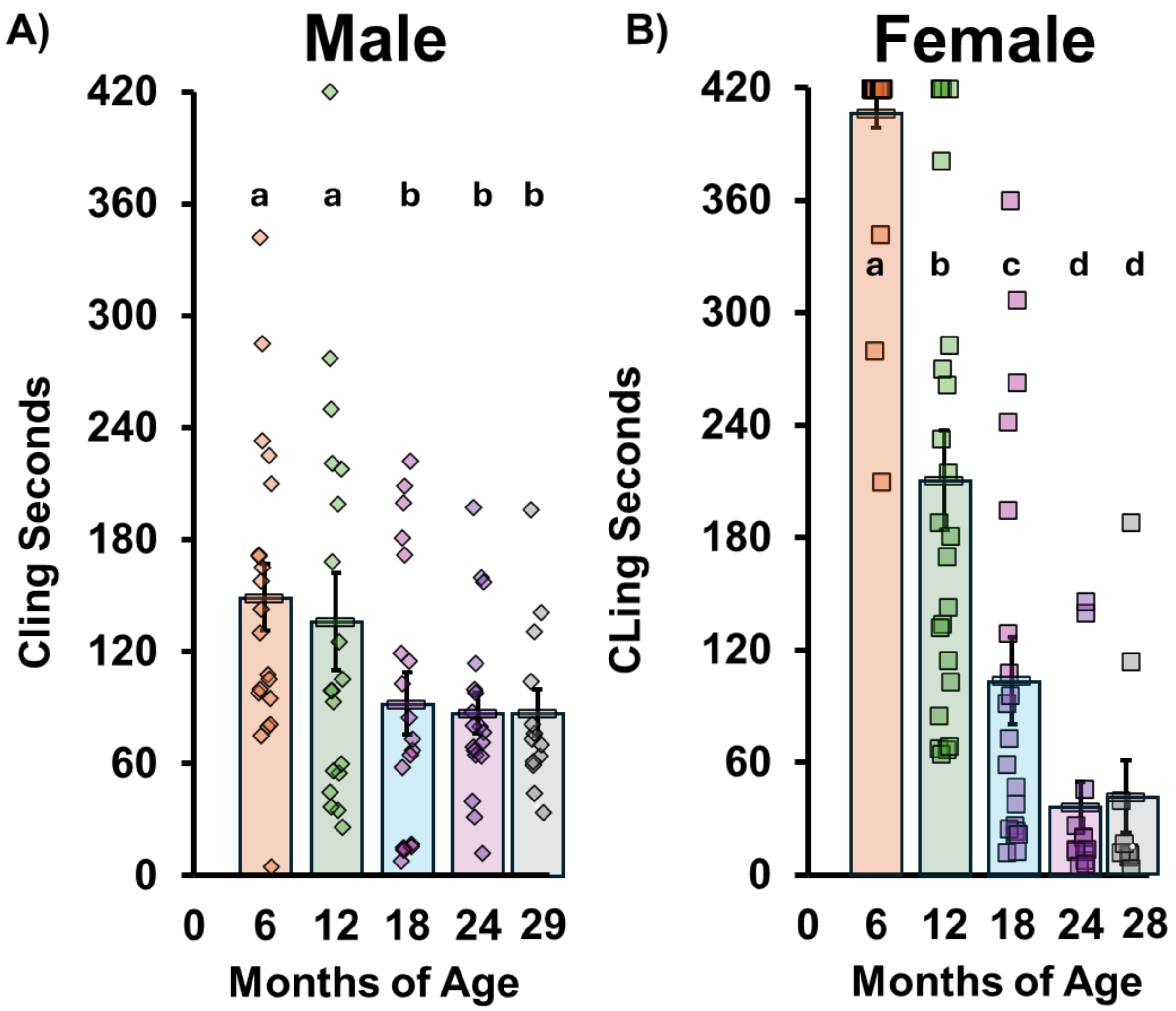


**Figure S5 Inverted Cling. A) Males B) Females. A) Males.) Females. Key:** m = meter, Each symbol (diamonds in Panel A and squares in Panel B) = the latency to fall in seconds of all the surviving mice at the given age. Different letters = significant difference (i.e., same letter = no difference in means), error bars = standard error, columns = mean.

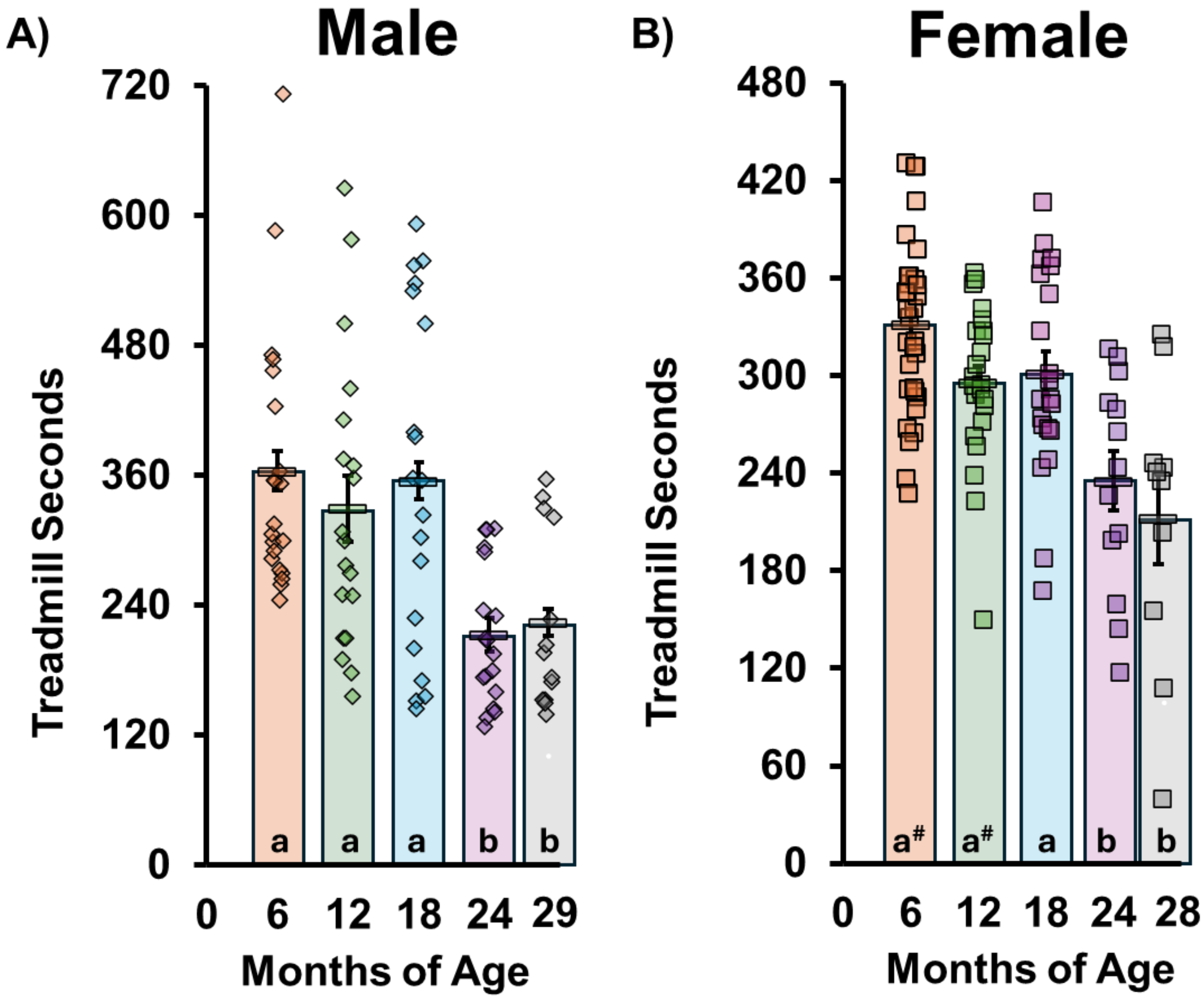


**Figure S6 Treadmill. A) Males B) Females. A) Males.) Females. Key**: m = meter, Each symbol (diamonds in Panel A and squares in Panel B) = the seconds to failure of all the surviving mice at the given age. Different letters = significant difference (i.e., same letter = no difference in means), # 0.05<p<0.10, error bars = standard error, columns = mean.

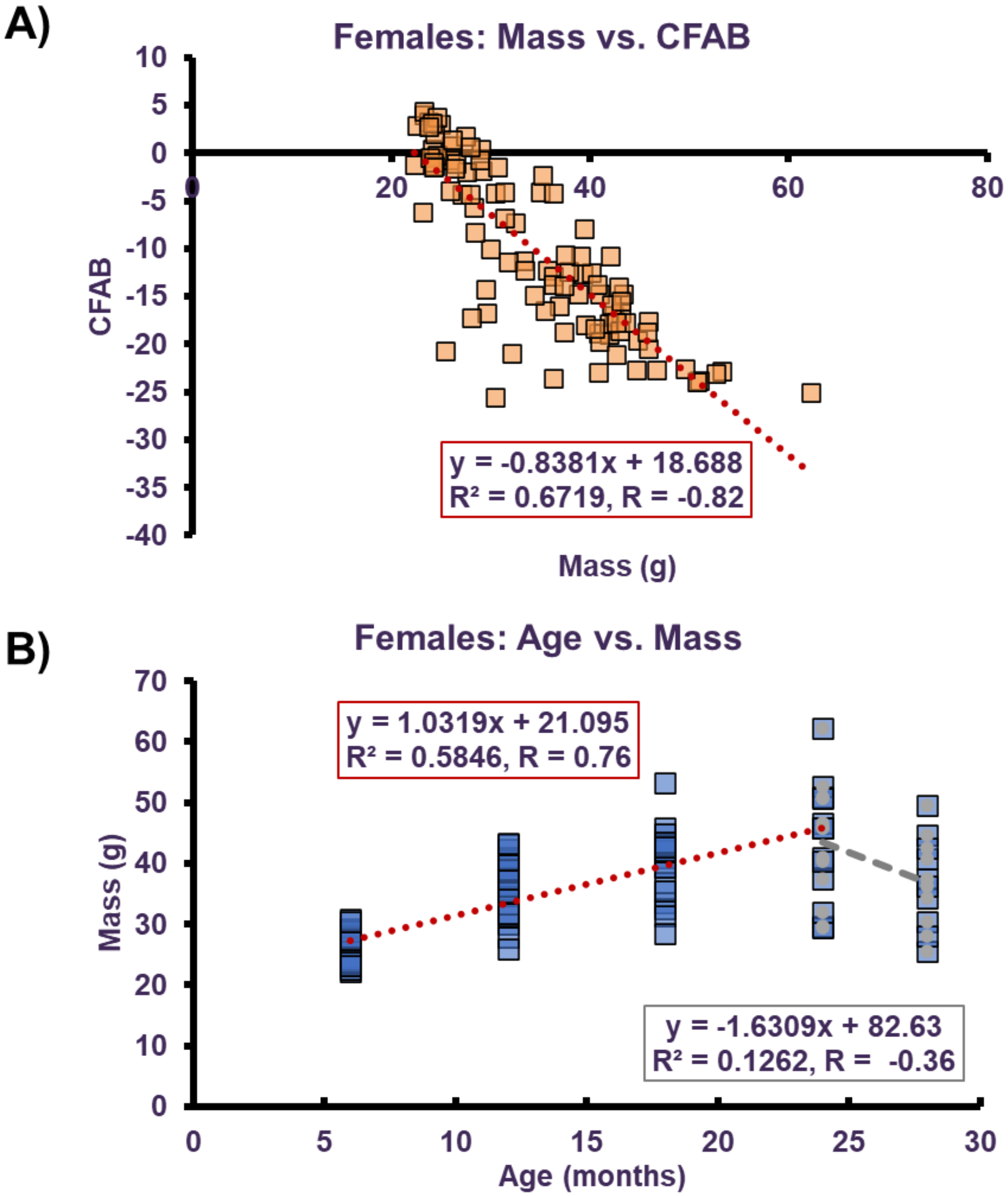


**Figure S7 Relationship between CFAB, Age and Body Mass in Females. A) Mass versus CFAB. B) Age versus Mass. Key:** m = meter, Each symbol (squares) = all the mice at each age in Panel A and all the surviving mice at the given age in Panel B. Equation is the simple linear regression.

**Supplemental Section References**
